# Reduced recombination relative to mutation characterizes dominant circulating clones of *Mycobacterium abscessus*

**DOI:** 10.64898/2026.09.03.749045

**Authors:** Chendi Zhu, Yu Zhou, Mingxing Ni, Zhuofan Huang, Zhenyu Wang, Junhao Zhu, Weimin Li

**Affiliations:** Beijing Chest Hospital, Capital Medical University, Beijing, 101149, China; Beijing Tuberculosis and Thoracic Tumor Research Institute, Beijing, 101149, China; Laboratory of Pathogen Microbiology and Immunology, Institute of Microbiology, Chinese Academy of Sciences, Beijing, China

**Keywords:** Mycobacterium abscessus, Nontuberculosis Mycobacteria, Circulating Clones, Evolution, Selective pressure

## Abstract

*Mycobacterium abscessus* exhibits extensive genomic diversity, yet independently emerged dominant circulating clones (DCCs) have achieved widespread distribution. However, the evolutionary changes associated with their emergence and subsequent diversification remain poorly understood. Here, we analyzed 11,314 genomes from 30 countries using a conservative framework integrating population-wide validation and read-supported sequence reconstruction, defining a stable core genome of 3,001 genes. Across seven DCCs, accessory-gene gains consistently exceeded losses on the ancestral branches leading to clone formation, with recurrent functions involving metal homeostasis, metabolism, environmental sensing and stress responses. Recombination also contributed substantial variation in specific lineages; in DCC3, ancestral recombinant regions encompassed core genes involved in iron acquisition, respiratory metabolism and protein homeostasis. Following DCC establishment, the relative contribution of homologous recombination to mutation consistently declined during within-clone diversification, indicating a broad shift toward mutation-dominated core-genome evolution. Together, these findings suggest a recurring pattern in DCC evolution, with accessory-genome gain and lineage-specific recombination occurring during clone formation, followed by increasingly mutation-dominated diversification after establishment.

## Introduction

*Mycobacterium abscessus* (Mab) is an increasingly important cause of difficult-to-treat pulmonary and extrapulmonary infections because of its extensive intrinsic antimicrobial resistance and poor treatment outcomes^1,2^. Although traditionally considered to be acquired independently from environmental reservoirs, population genomic studies have identified several closely related, globally distributed lineages termed dominant circulating clones (DCCs)^2-6^. Their emergence has challenged this conventional view and raised the possibility that particular lineages have evolved enhanced capacity for transmission, persistence, and adaptation to human hosts^3^.

Mab comprises three subspecies, *M. abscessus* subsp. *abscessus*, subsp. *massiliense*, and subsp. *bolletii*^5,7^, yet all seven currently recognized DCCs belong to the *abscessus* and *massiliense* subspecies^4^. In global collections, approximately 74% of individuals with cystic fibrosis are infected with DCC isolates, which have also been associated with enhanced virulence and poorer clinical outcomes^5,8^. These DCCs have undergone broadly synchronous population expansions and intercontinental dissemination^4^, and are also prevalent among individuals without cystic fibrosis^9^. However, why only a small subset of the diverse Mab population has evolved into globally successful circulating clones remains unclear.

Whole-genome sequencing has revealed a large, open Mab pangenome, with approximately 75% of a typical isolate’s genes belonging to the core genome^10,11^. Yet published core-genome estimates range from 3,354 to 4,159 genes^11-15^, reflecting differences in population diversity, analytical thresholds, annotation, and assembly quality and limiting comparisons across studies. Previous analyses of core-genome variation have nevertheless revealed substantial evolutionary diversity among Mab lineages and identified mutations associated with adaptation, antimicrobial resistance, and DCC evolution^2,3,9^. However, the presence of a gene across the population does not imply exclusively clonal inheritance of its sequence. Mab undergoes extensive homologous recombination, which can introduce sequence variation within core genes^12,16-18^. As recombinant variation is commonly removed from phylogenetic and evolutionary analyses, the contribution of recombination to core-gene evolution in Mab remains incompletely understood.

The accessory genome provides another major source of Mab diversity and has been implicated in DCC evolution^19^. Horizontal gene acquisition can alter genetic networks and generate lineage-specific phenotypes^20-22^ and functionally related genes, including global transcriptional regulators, have been independently acquired by several major DCCs^3^. DCC isolates also tend to have larger genomes but show reduced lateral gene transfer after establishment^12^. However, individual DCCs differ substantially in gene content, including phenylacetic acid metabolism in DCC1^15^, copper-associated mobile elements in DCC5^23^, and an *abscessus-*derived *rpoB* allele in DCC7^24^. Thus, although accessory-genome variation has been repeatedly associated with individual DCCs, whether DCC emergence follows a common pattern of accessory-genome evolution remains unclear.

Here, we analysed globally sampled Mab genomes using an integrated pangenome framework to examine the genomic changes accompanying DCC emergence and diversification. We defined a conservative core genome of 3,001 genes and found that DCC diversification was accompanied by a genome-wide shift toward mutation-dominated evolution, while recombination became concentrated in lineage-specific regions. Within DCC3, extensive ancestral recombination in one clade was concentrated in a core-genome region containing mycobactin biosynthesis and iron-acquisition genes, resulting in substantial recombinant sequence variation in these loci. Accessory-genome remodeling also occurred repeatedly across DCCs, with distinct gene-content changes converging on shared functional processes. Together, these findings reveal shared evolutionary patterns across independently emerged DCCs despite substantial lineage-specific genomic variation.

## Result

### Global pangenome analysis defines a population-representative core genome and reveals extensive gene-content diversity across DCCs

We assembled a global collection of 11,314 *Mycobacterium abscessus* (Mab) genomes from 30 countries (Fig. 1A, Supplementary Fig. 1A, Table S1). Sampling was biased towards Europe and North America, with Africa substantially underrepresented (Fig. 1A, Supplementary Fig. 1B). The dataset included 2,685 DCC1, 1,000 DCC2, 1,329 DCC3, 351 DCC4, 122 DCC5, 270 DCC6 and 195 DCC7 isolates. Treemmer-based^25^ stratified subsampling by phylogenetic diversity, country and subspecies yielded 1,130 representative genomes, including 793 DCC and 337 Non-DCC isolates, for pangenome analysis (Supplementary Fig. 1A, Table S2, Methods).

**Figure 1.**
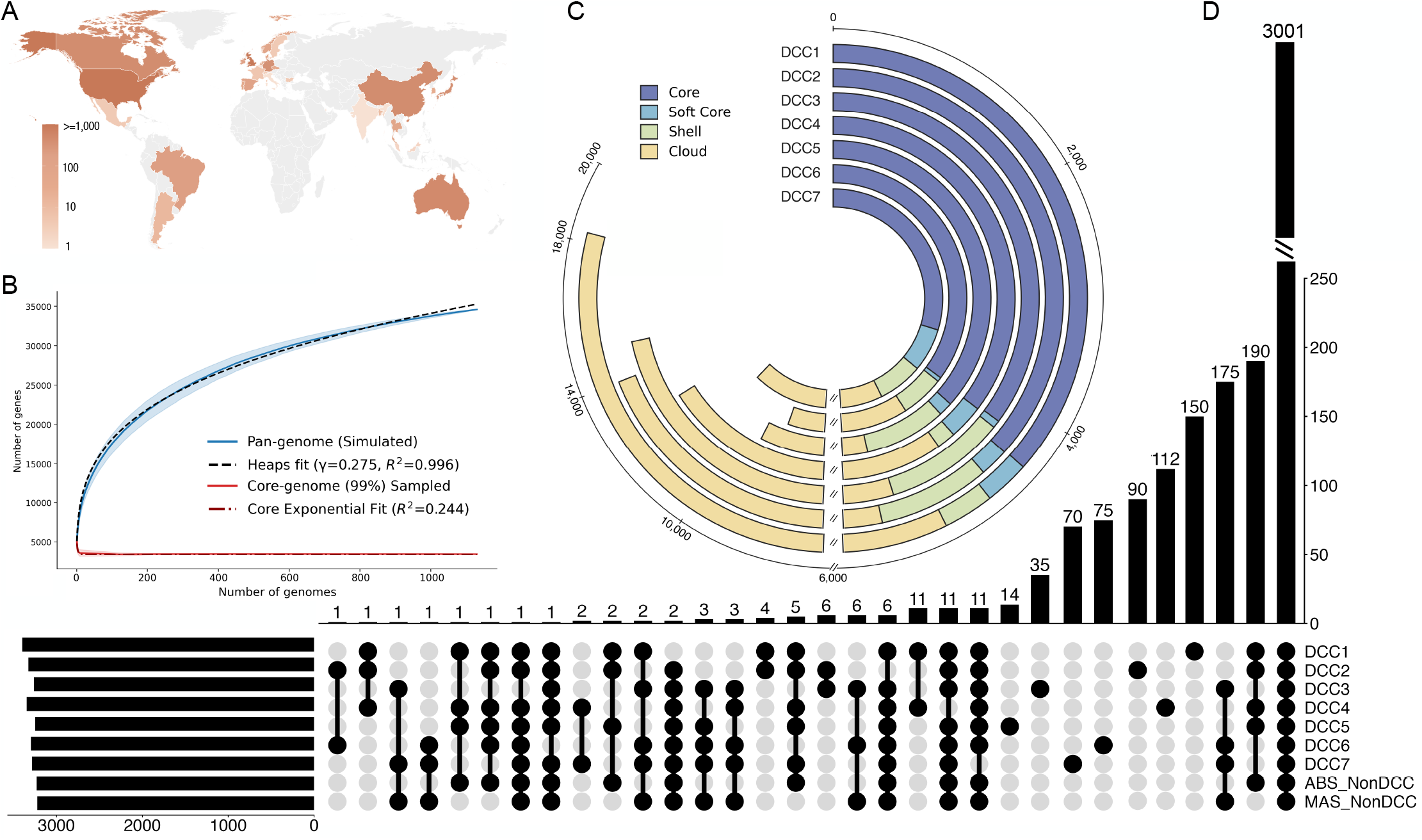
Global pangenome analysis defines a stable core genome of *Mycobacterium abscessus*. **(A)** Global distribution of *M. abscessus* genomes. Countries are shaded by isolate number on a logarithmic scale. **(B)** Pangenome and core-genome accumulation curves based on 1,130 diversity-preserving sampled genomes. Lines show the mean pangenome (blue) and 99%-prevalence core-genome (red) sizes across permutations, with shaded areas indicating variation. Heaps’ law fitting (γ = 0.275, R^2^ = 0.996) indicates an open pangenome. **(C)** Gene-frequency distributions across the seven DCCs. Bars indicate core (≥99%), soft-core (95–<99%), shell (15–<95%) and cloud (<15%) gene clusters. **(D)** UpSet plot of gene-cluster intersections among DCC1–DCC7 and non-DCC *abscessus* and *massiliense* populations. A total of 3,001 gene clusters were shared across all nine groups.

Pangenome analysis identified 34,571 gene clusters, including 3,436 core and 31,135 accessory clusters. The pangenome continued to expand with increasing sample size, with Heaps’ law fitting^26^ yielded γ = 0.275 (R^2^ = 0.996), consistent with an open Mab pangenome (Fig. 1B). We further examined whether this capacity for gene-content diversification persisted after the establishment of a dominant clone by independently analysing all 2,685 DCC1 isolates. DCC1 likewise retained an open pangenome (γ = 0.268, R^2^ = 0.978), suggesting that the capacity for accessory-gene acquisition was not substantially diminished following clonal establishment (Supplementary Fig. 1C).

The composition of the core genome, however, was strongly dependent on the population under consideration, consistent with previous pangenome studies. When each DCC was analysed independently, 3,562–4,401 genes were classified as core, consistently exceeding the 3,436 genes identified across the broader Mab population (Fig. 1C). Thus, genes that are accessory at the population level can become fixed or nearly fixed within individual DCCs, highlighting the importance of population representation in defining the Mab core genome. We therefore sought to establish a core-gene set that remained robust across the broader global population. The 3,436 genes initially classified as core by Panaroo^27^ were first aligned to the ATCC19977 reference genome to assess sequence completeness and alignment quality, resulting in the exclusion of 241 genes with >10% missing aligned sequence. The remaining 3,195 genes were subsequently evaluated across the remaining 10,184 genomes in the full dataset, and a further 194 genes present in <99% of isolates were removed. This filtering and population-wide validation defined a stable lower-bound core genome of 3,001 genes consistently represented across the currently available Mab population (Fig. 1D, Supplementary Fig. 1A, Table S3).

Having established this population-wide core genome, we next examined whether independently emerged DCCs shared common accessory genes. Group-associated genes were defined as those present in >90% of isolates within a target group but ≤10% outside that group. Most associations were specific to individual DCCs or reflected subspecies structure, with only a small number shared among subsets of DCCs. No gene cluster was shared by all seven DCCs while simultaneously absent from Non-DCC populations of both subsp. *abscessus* and subsp. *massiliense* (Fig. 1D, Supplementary Fig. 1D). These findings suggest that the independent emergence of DCCs is unlikely to be driven by a common set of accessory genes.

### Read-supported core-genome analysis refines the phylogenetic structure of DCCs

Having defined a stable set of 3,001 core genes, we next evaluated whether SNPs inferred from assembly-based alignments were supported by the underlying sequencing reads. Assembly-derived SNPs were compared with consensus calls generated by direct read mapping to the ATCC 19977 reference genome. These were classified as “supported” when both approaches identified the same alternative allele, “inconsistent” when they produced different allele states, or “missing” when read depth or allele frequency was insufficient for a confident mapping-based call (Fig. 2A).

**Figure 2.**
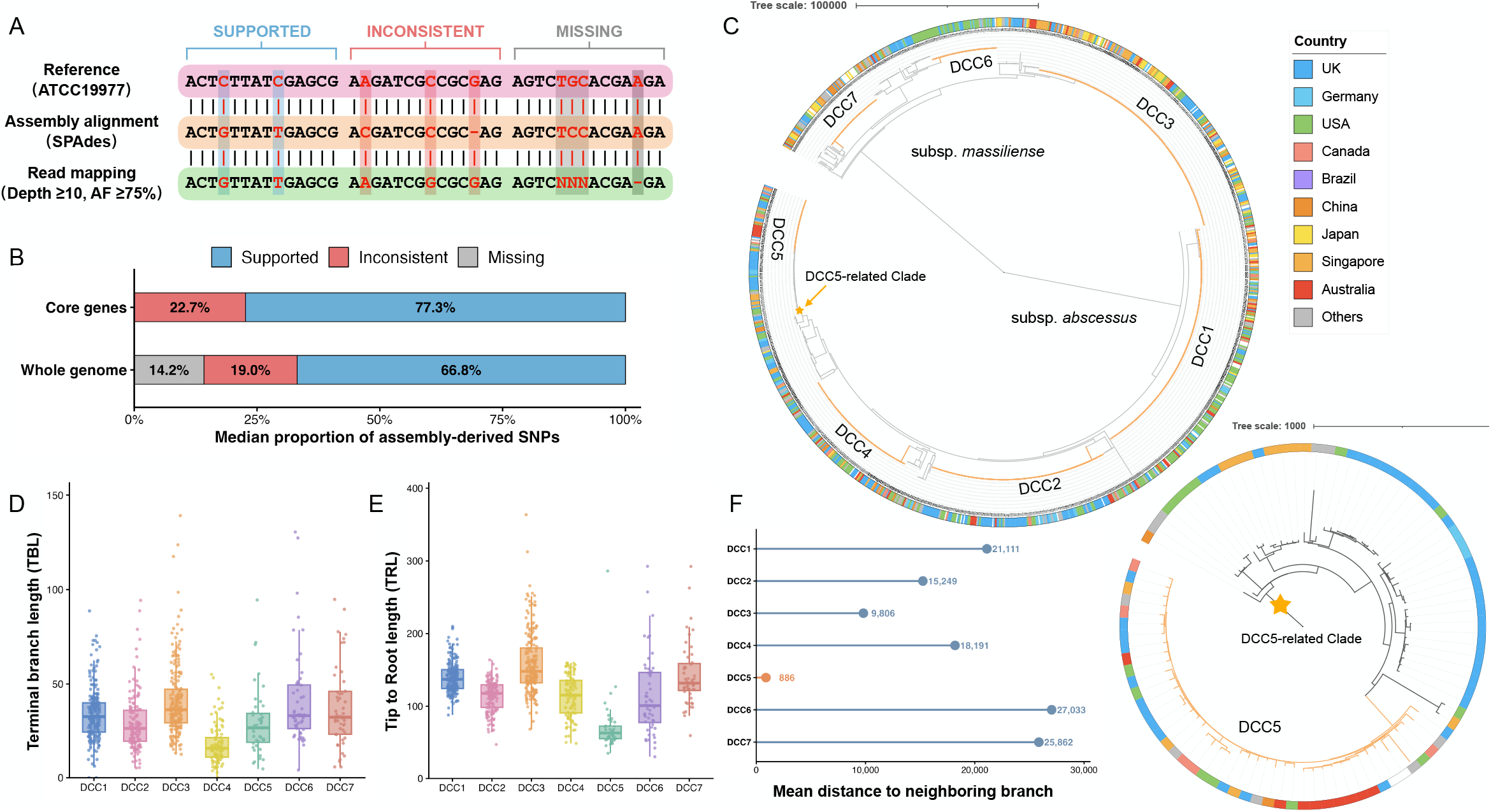
Construction of a stable, read-supported core-genome alignment. **(A)** Comparison of SNPs inferred from SPAdes assemblies and direct read mapping to the ATCC 19977 reference. Assembly-derived SNPs were classified as supported, inconsistent or missing based on mapping consensus. Mapping calls required ≥10× depth and ≥75% alternative allele frequency. **(B)** Median proportions of supported, inconsistent and missing assembly-derived SNPs across the whole genome and stable core genome. **(C)** Maximum-likelihood phylogeny reconstructed from the read-supported core-genome alignment after masking recombination-associated regions identified by Gubbins. The outer ring indicates country of isolation, and orange arcs denote DCCs. The inset shows the phylogenetic position of DCC5. **(D)** Distribution of terminal branch lengths (TBLs) across DCCs. **(E)** Distribution of tip-to-root lengths (TRLs) across DCCs. **(F)** Mean pairwise genetic distances between each DCC and its neighboring phylogenetic lineage.

Across the whole genome, a median of 66.8% of assembly-derived SNPs were supported by read mapping, while 19.0% were inconsistent and 14.2% lacked confident mapping calls (Fig. 2B, Supplementary Fig. 2A, Table S4). Restriction to the stable core genome largely eliminated missing calls, but 22.7% of assembly-derived core SNPs remained inconsistent with the mapping consensus. These discrepancies were broadly distributed across the 3,001 core genes rather than concentrated in a small number of problematic loci (Supplementary Fig. 2B), indicating that locus-specific exclusion would not resolve the discrepancy. We therefore adopted a conservative read-supported strategy, retaining only nucleotide states meeting predefined read-depth and allele-frequency thresholds and treating the remaining positions as missing. The resulting reference-coordinate core-genome alignment was used for subsequent analyses.

After masking recombination-associated regions, the resulting maximum-likelihood phylogeny preserved the overall DCC structure, with individual DCCs forming compact phylogenetic groups (Fig. 2C). Terminal branch lengths (TBLs) and tip-to-root lengths (TRLs) retained substantial within-DCC variation, indicating preservation of their internal genetic diversity (Fig. 2D, E). DCC5 was an exception, being nested within a broader closely related lineage rather than separated by a long ancestral branch (Fig. 2C). Mean within-DCC5 divergence was 71.5 SNPs compared with 886 SNPs between DCC5 and the neighboring lineage, supporting its genetic distinction, although this separation was substantially smaller than for other DCCs (9,806–27,033 SNPs; Fig. 2F). Similar nesting of DCC5 has been observed in previous global Mab phylogenies^9^, suggesting that DCC5 represents a less phylogenetically isolated DCC whose apparent boundary may vary with the treatment of recombination-associated variation.

### Core-genome evolution shifts toward mutation-dominated diversification during DCC expansion

We next investigated how the relative contributions of mutation and homologous recombination changed during DCC evolution. For each DCC, the branch leading to its most recent common ancestor represented the ancestral phase, whereas descendant branches captured diversification following clone establishment (Fig. 3A). Recombination- and mutation-derived SNPs were assigned to phylogenetic branches and mapped to the 3,001 stable core genes, allowing gene-specific changes in the recombination-to-mutation ratio (r/m) to be compared across this transition.

**Figure 3.**
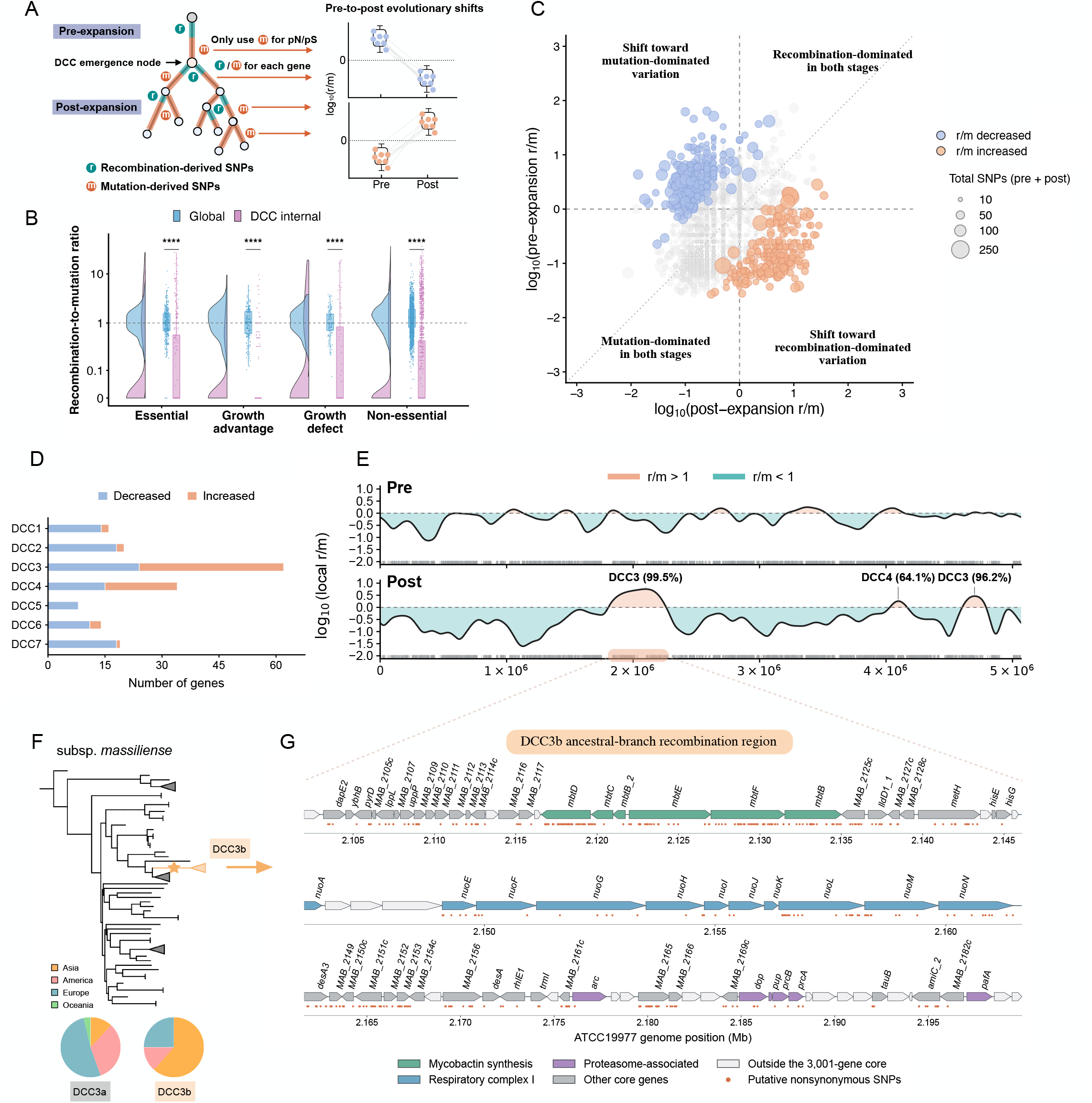
Global reduction and localized retention of homologous recombination during DCC diversification. **(A)** Branch-based framework comparing evolutionary processes before and after DCC diversification. The stem branch represents the ancestral phase, and descendant branches represent subsequent diversification. r/m ratios were calculated from recombination- and mutation-derived SNPs. **(B)** Distribution of gene-specific r/m ratios across the global population and individual DCCs, stratified by transposon-sequencing gene classes. Dashed line indicates r/m = 1. **(C)** Pre- and post-diversification r/m comparison across core genes. Blue and orange indicate genes with decreased (n = 288) and increased (n = 263) r/m, respectively. Point size indicates total SNP events. **(D)** Numbers of genes showing decreased or increased r/m within each DCC among genes identified in panel C. **(E)** Genome-wide landscape of r/m variation across the 3,001 core genes. Curves represent smoothed recombinant-to-mutation SNP ratios across the ATCC19977 reference genome. Highlighted peaks indicate regions with elevated recombination contribution and the dominant contributing DCC. **(F)** Phylogenetic localization of the DCC3b ancestral branch associated with the major recombination peak. **(G)** Gene organization of the DCC3b-associated recombination region near 2.1 Mb. Colors indicate functional categories, including mycobactin synthesis, respiratory complex I, proteasome-associated genes, and other core genes. Orange dots indicate recombinant SNP positions.

Core genes were further classified as essential, growth-defect, growth-advantage or non-essential based on multi-strain TnSeq data^14^. Across all four classes, including essential genes, r/m was consistently and substantially lower within DCCs than in the broader Mab population (Fig. 3B, Table S5), demonstrating that the reduced contribution of recombination extended across functionally distinct components of the core genome. Overall, this pattern was consistent with a broad shift toward mutation-dominated clonal diversification in DCCs. Notably, 48 TnSeq-defined essential genes were absent from the stable core set. Of these, 18 were excluded because of incomplete alignments, while the remainder showed substantial sequence divergence between subspecies, preferential presence in subsp. abscessus, or absence from subsets of isolates (Supplementary Fig. 3A, Table S6). Thus, functional essentiality and population-wide genomic conservation were not fully concordant.

Despite the genome-wide decline in the relative contribution of recombination, 263 core genes showed significantly increased r/m following DCC expansion (Fig. 3C). These genes spanned all four TnSeq-defined functional classes and were enriched in respiratory metabolism, oxidative phosphorylation, proteasome function and other metabolic pathways (Supplementary Fig. 4A–D). To determine the origin of these localized increases, we reconstructed cumulative recombination- and mutation-derived variation along the phylogenetic expansion of each DCC (Supplementary Fig. 4B). Although all seven DCCs showed an overall pattern in which mutation accumulated more rapidly than recombination after expansion, DCC3 and DCC4 showed transient increases in recombination during specific stages of their diversification. Consistent with this pattern, DCC3 and DCC4 contained substantially more genes with increased r/m than the other DCCs (Fig. 3D).

We therefore mapped changes in r/m across the genome to determine whether these increases reflected dispersed gene-specific effects or localized recombination events. Genes with elevated r/m were strongly clustered rather than randomly distributed, forming prominent peaks near 2.1, 4.1 and 4.7 Mb (Fig. 3E). The peaks at approximately 2.1 and 4.7 Mb were predominantly attributable to DCC3. Tracing the DCC3-associated recombinant variation back through the phylogeny further localized these signals to a secondary ancestral node within DCC3 (Fig. 3F). This node marked a major internal subdivision of DCC3 corresponding to the previously described DCC3a and DCC3b sublineages^28^, indicating that much of the apparent gene-level increase in r/m resulted from ancestral recombination associated with diversification of an internal DCC3 lineage.

We next examined the recombinant regions associated with the ancestral branch leading to DCC3b. The major peak near 2.1 Mb comprised an extended recombinant region encompassing approximately 63 core genes and showed clear functional organization (Fig. 3G). Three major functional blocks were evident: the mycobactin biosynthesis locus, including multiple *mbt* genes involved in siderophore-mediated iron acquisition; respiratory complex I genes involved in energy metabolism and redox homeostasis; and the *Pup*-proteasome-associated region, including *dop, prcA/prcB* and related factors involved in regulated protein degradation and stress responses. DCC3a and DCC3b also differed in their geographic composition: 59.9% (91/152) of DCC3b isolates originated from Asia, compared with 11.7% (120/1,024) of DCC3a isolates, indicating substantial geographic structuring between the two sublineages (Fig. 3F).

Together, these analyses show that the increased r/m observed at a subset of core genes was largely attributable to localized ancestral recombination within specific DCC lineages rather than a reversal of the broader evolutionary trend. Thus, the overall decline in r/m remained the predominant pattern during DCC diversification, while the increased r/m observed in a subset of genes largely reflected localized ancestral recombination events in specific DCC lineages.

### Recombination and selective constraints reflect distinct dimensions of DCC evolution

We next compared changes in recombination with previously identified shifts in selective constraint during DCC evolution^29^. Within the stable core genome, 48 genes showed high-confidence shifts in selective regime, involving functions related to cell-envelope remodeling, lipid metabolism, stress responses and metal homeostasis (Supplementary Fig. 5A, B; Tables S7, S8). These genes showed limited overlap with loci exhibiting altered r/m, most genes with selective-regime shifts showed no corresponding change in recombination contribution, while most genes with altered r/m did not cross the pN/pS = 1 threshold (Supplementary Fig. 4E). Thus, changes in recombination and selective constraint were largely uncoupled at the gene level.

DCC5 showed a distinct pattern of selective evolution, lacking many of the recurrent selective-regime shifts observed across the other DCCs (Supplementary Fig. 5C). Given its shorter ancestral separation and nested phylogenetic position, we further examined whether DCC5 had already acquired variants in recurrently selected genes, including *phoR, ideR* and *ubiA*, before its diversification. No such pattern was observed (Supplementary Fig. 5D), indicating that the absence of recurrent selective shifts in DCC5 could not be explained by prior acquisition of these variants. Overall, shifts in selective constraint were predominantly gene- and lineage-specific and were largely distinct from the broader change in the mutation–recombination balance accompanying DCC diversification.

### Convergent accessory-genome expansion accompanies DCC emergence

Although no accessory gene was universally associated with all DCCs, we next examined whether broader patterns of accessory-genome evolution accompanied DCC emergence. Using the complete accessory-gene presence–absence matrix, the resulting population structure closely recapitulated the stable core-genome phylogeny, recovering the major subspecies division and maintaining individual DCCs as distinct clusters (Fig. 4A, Supplementary Fig. 6A, B). Thus, despite extensive variation in gene content, accessory-genome structure remained closely associated with lineage history.

**Figure 4.**
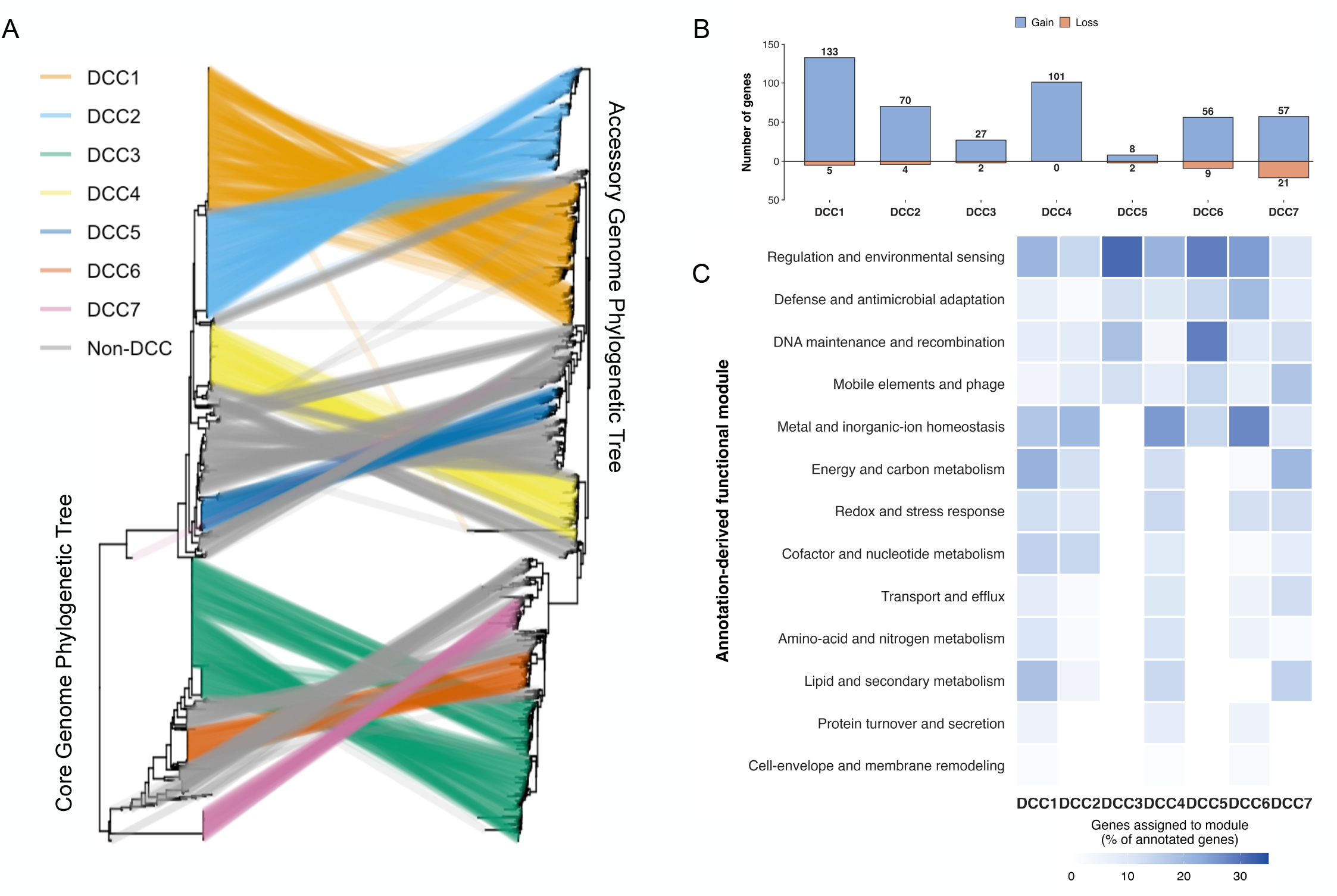
Accessory-genome evolution during DCC emergence is dominated by lineage-specific gene gain. **(A)** Comparison of phylogenetic structures reconstructed from the stable core genome and accessory-gene presence–absence matrix. Lines connect identical isolates between trees and are colored by DCC assignment; non-DCC isolates are shown in grey. **(B)** Accessory-gene gain and loss events reconstructed on the stem branch leading to each DCC. Blue and orange bars indicate gene gains and losses, respectively, with values showing the number of inferred events. **(C)** Functional composition of accessory genes gained on DCC stem branches. eggNOG annotations were grouped into broader functional modules, with colors indicating the proportion of annotated gained genes assigned to each module within each DCC. White indicates no genes assigned to the corresponding module.

We next reconstructed accessory-gene gains and losses along the ancestral branch leading to each DCC. Gene gains consistently exceeded losses across all seven DCCs, with 8–133 gains compared with 0–21 losses (Fig. 4B, Table S9), revealing a shared pattern of gain-dominated accessory-genome expansion during DCC emergence. Most gained genes showed taxonomic affinities within *Mycobacterium sensu lato*, with additional matches to other actinobacterial genera, including *Nocardia, Gordonia* and *Streptomyces* (Supplementary Fig. 6C, D). DCC1 showed the strongest affinity to *M. abscessus*, consistent with the extensive representation of the DCC1 reference strain ATCC19977 in current annotation databases. Because these assignments were based on eggNOG seed orthologs, they were used to describe taxonomic affinities rather than infer direct gene origins.

Despite lineage-specific differences in the genes gained, their functions showed partial convergence across DCCs (Fig. 4C, Table S9). Recurrently represented functions included regulation and environmental sensing, metal and inorganic-ion homeostasis, energy and carbon metabolism, redox and stress responses, and DNA maintenance and recombination, whereas mobile elements and phage-associated functions were more lineage restricted. Thus, accessory-genome expansion during DCC emergence involved different genes but recurrently affected similar functional categories.

## Discussion

In this study, we identified shared genome-wide evolutionary patterns accompanying the emergence and diversification of seven *Mycobacterium abscessus* DCCs. Using a conservative core-genome framework, we found a consistent decline in recombination relative to mutation following DCC expansion. This genome-wide trend was accompanied by localized ancestral recombination affecting discrete core-genome regions in specific lineages. Accessory-gene gains also consistently exceeded losses during DCC formation, with different genes recurrently affecting similar functional categories. Together, these findings suggest that independently emerged DCCs share broad evolutionary patterns despite substantial lineage-specific genomic variation.

Defining a reliable core genome remains challenging for Mab because estimates are strongly influenced by population sampling and genome quality^11-15^. Most available Mab genomes are reconstructed from short reads, for which assembly errors can generate false-positive variants^30^, inflate apparent recombination, and produce artificial gene absence^31^. We therefore combined phylogenetically representative sampling, gene-level quality control, population-wide validation, and read-supported nucleotide reconstruction to define a conservative set of 3,001 core genes. This set should be regarded as a stable lower bound for the currently sampled Mab population rather than a biological definition of the core genome. This distinction is reinforced by Tn-seq studies showing that experimentally essential genes are not necessarily universally conserved and that essentiality can vary among genetic backgrounds^14,32,33^. Long-read sequencing can resolve many of these limitations, but complete Mab genomes remain limited and current studies generally include only a few to several dozen isolates^13,34,35^. Our framework therefore provides a practical basis for population-scale evolutionary analysis with existing short-read data, while the genomic boundaries of the Mab core will continue to be refined as population-scale long-read sequencing becomes available.

Previous studies have reported reduced lateral genetic exchange in Mab DCCs relative to environmentally acquired isolates, often interpreted as a feature associated with host adaptation^12^. However, these comparisons generally treated established DCC populations collectively^3,4,12^, making it difficult to determine when this reduction emerged. By resolving the seven DCCs separately and tracing recombination and mutation along their phylogenetic expansion, we found that declining r/m was a recurrent transition accompanying DCC diversification rather than simply a static property of DCC membership. Its consistency across all TnSeq functional classes, including essential genes, further indicates that this transition extends broadly across the core genome. The resulting predominance of mutation over recombination is consistent with increasingly vertical, clonal diversification after DCC establishment.

Importantly, this genome-wide decline did not eliminate the evolutionary contribution of recombination. A subset of core genes retained increased r/m, particularly those involved in respiratory metabolism, oxidative phosphorylation, proteasome function, and other metabolic processes. Our phylogenetic reconstruction showed that many of these gene-level increases could be traced to localized ancestral recombination events in specific lineages, particularly DCC3. In DCC3, a major recombinant region on the ancestral branch leading to DCC3b encompassed functionally relevant modules involved in iron acquisition, respiratory metabolism and protein homeostasis. The region included the mycobactin biosynthesis locus, which is required for growth of Mab under iron limitation and contributes to intracellular replication in macrophages^36,37^. Studies in *M. tuberculosis* similarly demonstrate an essential role of mycobactin-mediated iron acquisition during iron limitation and intracellular growth^38,39^. The same region also contained respiratory complex I genes involved in energy production and redox homeostasis, pathways that are extensively remodeled in Mab under nutrient and metal stress^40,41^, as well as components of the *Pup*-proteasome system, including *dop* and *prcA/prcB*^42^. In *M. tuberculosis*, this system contributes to protein quality control, stress resistance and persistence during chronic infection^43^. Thus, this ancestral recombination event generated extensive allelic variation across core genes involved in iron acquisition, bioenergetics and stress responses. Together with the geographic enrichment of DCC3b in Asia, these findings raise the possibility that ancestral recombination contributed to lineage-specific adaptation, although the functional effects of these recombinant alleles remain to be determined.

Changes in selective constraint provide a related but largely distinct view of this diversification. The limited overlap between genes with altered r/m and those showing shifts in pN/pS indicates that changes in recombination were not generally accompanied by corresponding changes in selective regime. Whereas declining r/m represents a broad shift in the relative sources of core-genome variation, pN/pS changes were predominantly gene- and lineage-specific. Recurrent shifts involving regulators such as *phoR, ideR*, and *ubiA* nevertheless suggest that some functional systems continue to experience changing selective pressures during clonal diversification^3,29^. DCC5 was an exception to several of these recurrent patterns, potentially reflecting its nested phylogenetic position and less discrete ancestral boundary. Thus, selection analyses support continued lineage-specific adaptation but do not account for the broader decline in recombination that characterizes DCC expansion.

Accessory-genome evolution suggests that a different pattern predominated during DCC formation. Mab has a highly dynamic accessory genome shaped by horizontal gene transfer, and previous studies have proposed that accessory-gene acquisition contributed to the emergence of major DCCs^12,18,44^. Across all seven DCCs, we found that gene gains exceeded losses along their ancestral branches, but no acquired gene was universally shared. Instead, independently acquired genes recurrently involved environmental sensing, metabolism, metal homeostasis, and stress responses. This combination of lineage-specific gene acquisition and functional convergence suggests that DCC formation did not depend on a common set of accessory genes, but repeatedly involved remodeling of related biological functions. Together with the subsequent genome-wide decline in r/m, these findings are consistent with a transition from greater genome remodeling around clone formation toward predominantly vertical diversification after establishment.

Several limitations affect the interpretation of these patterns. Publicly available genomes are unevenly distributed across geographic regions, studies and patient populations, which may influence both lineage frequencies and geographic associations. The stem branches used here provide an operational definition of ancestral change but do not directly identify the timing of ecological transitions or population expansion. Estimates of r/m describe the relative contribution of inferred recombination to sequence variation and do not measure the absolute rate of genetic exchange; they may also depend on donor divergence, mapping sensitivity and phylogenetic sampling. Finally, functional annotations identify candidate biological processes affected by gene acquisition or recombinant alleles, but their effects on bacterial fitness, host adaptation and transmission remain to be established experimentally.

In conclusion, our study reveals a consistent shift in genome evolution accompanying the emergence and expansion of Mab DCCs. DCC formation was characterized by gain-dominated accessory-genome remodeling, whereas subsequent expansion was accompanied by a reduced contribution of recombination and reorganization of selective pressures in the core genome. Despite substantial variation in the specific genes involved, convergence emerged at the level of evolutionary processes and functional programs, indicating that successful circulating clones may follow broadly similar evolutionary trajectories through distinct genetic and ecological routes. These findings provide a genome-wide perspective on how Mab DCCs emerge and diversify and help reconcile their shared evolutionary features with the substantial heterogeneity observed among individual clones.

## Materials and Methods

### Genome sequence collection

We systematically searched PubMed and the NCBI Sequence Read Archive (SRA) up to 1 March 2025 to identify publicly available whole-genome sequencing data for *Mycobacterium abscessus* (Mab). Searches were performed using the terms “*Mycobacterium abscessus*” and “*Mycobacteroides abscessus*”. For studies identified through PubMed, the associated BioProject and sequencing-run accessions were retrieved from the original articles, supplementary materials, or linked NCBI records. In total, 188 BioProjects comprising 11,702 putative Mab whole-genome sequencing samples were identified. Following species confirmation, removal of low-quality or contaminated datasets, and deduplication, 11,314 nonredundant Mab isolates were retained for downstream analyses (Table S1). These isolates originated from 30 countries or regions across Asia, Europe, Oceania, and the Americas.

### De novo assembly and subspecies assignment

Raw sequencing reads were assembled de novo using SPAdes v3.11.1^45^ with the --careful option and a PHRED quality offset of 33 (--phred-offset 33). The resulting assemblies were compared with representative genomes of the three recognized Mab subspecies: *M. abscessus* subsp. *abscessus* ATCC 19977 (NC_010397.1), subsp. *massiliense* CCUG 48898 (NZ_AP014547.1), and subsp. *bolletii* GD91 (NZ_CP065265.1).

Genome-wide average nucleotide identity was calculated using *fastANI* (v1.2)^46^. Each isolate was assigned to the subspecies represented by its highest-scoring reference genome, provided that the corresponding ANI was at least 98%. All 11,314 retained isolates met this criterion and were assigned to one of the three subspecies (Table S1).

### DCC phylogenetic reconstruction and diversity-preserving sampling

To obtain a phylogenetically representative dataset while limiting redundancy among densely sampled dominant circulating clones (DCCs), phylogenetic reconstruction and subsampling were performed separately for each DCC. Draft genomes generated by SPAdes v3.11.1^45^ were placed in a common reference-coordinate alignment. Putative recombinant regions were identified and masked using Gubbins^47^, and maximum-likelihood phylogenies were reconstructed from the resulting recombination-filtered alignments. The complete, unsampled DCC-specific trees were used for diversity-preserving isolate selection. Phylogeny trees were visualized in *FigTree* (v1.4.4)^48^ or *iTOL*^*49*^.

TreeMmer^25^ was applied independently to each DCC phylogeny. TreeMmer^25^ iteratively removes terminal isolates that contribute the least unique branch length, thereby reducing sample size while retaining a predefined proportion of the phylogenetic diversity represented by the original tree. To balance unequal DCC sample sizes against the retention of rare lineages, the retained tree-diversity targets were 50% for DCC1, 70% for DCC2–DCC4, and 90% for DCC5–DCC7.

Non-DCC isolates were sampled separately by stratifying the dataset according to subspecies and country or region, thereby maintaining representation of the principal taxonomic and geographic strata. This procedure yielded a representative dataset of 1,130 genomes, comprising 793 DCC and 337 Non-DCC isolates, for pangenome reconstruction.

### Pangenome reconstruction

The 1,130 representative assemblies were annotated using Prokka^50^. The resulting GFF files were analyzed using Panaroo^27^ in strict cleaning mode to reduce gene clusters arising from annotation errors, fragmented assemblies, and other assembly-associated artifacts. Paralog splitting was enabled to improve discrimination among duplicated homologues. Panaroo^27^ identified 34,571 nonredundant gene clusters. Genes detected in at least 99% of the representative genomes were provisionally classified as core genes, yielding an initial core genome of 3,436 genes. The remaining 31,135 gene clusters were classified as components of the accessory genome.

### Alignment-quality filtering and independent validation of the stable core genome

Because frequency-based core-genome definitions may retain fragmented or poorly aligned loci, each Panaroo^27^ core-gene alignment was subjected to an additional alignment-quality assessment. For each alignment column, the proportion of sequences containing a gap was calculated. Columns containing gaps in more than 50% of sequences were classified as gap-dominated. A gene was retained only when gap-dominated columns accounted for no more than 10% of its alignment length. This filtering procedure removed 241 poorly aligned loci and retained 3,195 quality-controlled core genes. The prevalence of these genes was subsequently re-evaluated in the isolates that were not included in the representative pangenome dataset. Genes detected in more than 99% of this independent validation set were retained, producing a final stable core-genome set of 3,001 genes. Because short-read assembly, gene fragmentation, and annotation failure can cause genuine core genes to appear absent or poorly aligned, the resulting 3,001-gene set was interpreted as a conservative lower bound on the stable Mab core genome rather than an exhaustive catalogue of all biologically core loci.

### Identification of DCC-associated and group-specific genes

Custom Python scripts were developed to extract DCC-associated genes from the complete Panaroo^27^ gene presence–absence matrix (*gene_presence_absence*.*Rtab*). Isolate identifiers in the matrix were first matched to their corresponding DCC or Non-DCC assignments. For each gene cluster, the carriage frequency was calculated independently within DCC1–DCC7 and the two subspecies-matched Non-DCC populations. Gene clusters were then summarized according to all groups in which they were detected, generating a group-combination profile for each gene.

For each DCC, the accessory-gene list included all non-core gene clusters carried by that DCC whose group-combination profile did not include a Non-DCC population. Genes detected in more than one DCC but absent from Non-DCC isolates were retained in the gene list of every corresponding DCC and were additionally classified as shared DCC-associated genes. Genes detected exclusively in a single DCC were classified as DCC-specific genes. This combination-based approach allowed DCC-restricted genes to be distinguished from accessory genes broadly distributed in the background Mab population without imposing an additional prevalence threshold during this classification step.

A complementary frequency-based screen was performed using a custom script to identify strongly group-specific gene clusters. For each focal DCC, genes were considered enriched when their carriage frequency was greater than 90% within the focal group and less than 10% among isolates outside that group. Conversely, putative DCC-associated gene losses were defined as genes carried by less than 10% of isolates within the focal DCC but by more than 90% of isolates outside that DCC. The scripts reported the number and percentage of gene-positive isolates in every group, enabling all candidate gains, losses, and DCC-specific genes to be inspected against their complete population-level distribution.

### SNP calling

Raw sequencing reads were quality-trimmed using Sickle^51^, retaining bases with PHRED quality scores greater than 20 and reads longer than 30 bp. The filtered reads from all isolates, irrespective of subspecies, were mapped to the *M. abscessus* subsp. *abscessus* ATCC 19977 reference genome (NC_010397.1) using BWA-MEM v0.7.17^52^. Using a single reference genome provided a common genomic coordinate system for variant comparison and phylogenetic reconstruction across the complete Mab population. Mapping files were sorted and indexed using SAMtools v1.3.1^53^. Variants were initially called using SAMtools^53^ and VarScan2 v2.3.9^*54*^. Sites supported by a sequencing depth of at least 20 reads were retained. Variants with an allele frequency of at least 75% were classified as fixed mutations. Small insertions and deletions were identified concurrently using VarScan2^54^. Variants located within annotated repetitive or potentially ambiguously mapped regions, including prophages, insertion sequences, and mobile genetic elements, were excluded from downstream SNP analyses.

### Comparison of assembly-derived and read-supported SNPs

To assess the reliability of SNPs inferred from draft assemblies, we compared assembly-derived whole-genome alignments with consensus sequences generated directly from read mapping. SPAdes^45^ assemblies were aligned against the ATCC 19977 reference genome to generate a reference-coordinate assembly alignment. Independently, quality-filtered sequencing reads from the same isolates were mapped to ATCC 19977 using BWA-MEM^52^.

A genomic position was considered confidently callable from read mapping when it was covered by at least 10 reads and the predominant allele had a frequency of at least 75%. At each variable position identified in the assembly alignment, the assembly-derived allele was compared with the corresponding read-supported consensus allele. Assembly-derived SNPs were classified as supported when the assembly and read-mapping alleles were identical, inconsistent when the site was confidently callable but the predominant read-supported allele differed from the assembly-derived allele, and missing when the read-mapping data did not meet the depth or allele-frequency thresholds.

The proportions of supported, inconsistent, and missing assembly-derived SNPs were calculated for each isolate, first across the whole reference-aligned genome and subsequently within the 3,001-gene stable core genome. Problematic sites within the core genome were assigned to genes according to their ATCC 19977 coordinates to determine whether assembly–mapping inconsistencies were concentrated in particular loci.

Because a substantial proportion of assembly-derived SNPs was not supported by direct read mapping, the final alignment used for recombination inference was reconstructed from mapped reads. For each isolate, a full-length consensus sequence was generated according to the ATCC 19977 coordinate system. At positions with a sequencing depth of at least 10 and a predominant allele frequency of at least 75%, the predominant nucleotide was inserted into the consensus sequence. Positions that failed either threshold were represented as N, rather than being imputed as the ATCC 19977 reference allele.

The consensus sequences of all isolates were combined into a common reference-coordinate multiple-sequence alignment. Positions containing unresolved nucleotides in more than 5% of isolates were excluded before phylogenetic reconstruction. This procedure retained confidently supported reference and alternative alleles while reducing false SNPs caused by assembly errors, inadequate read coverage, or ambiguous mapping.

### Terminal branch lengths (TBL) and tip-to-root lengths (TRL)

Terminal branch length (TBL) and tip-to-root length (TRL) were calculated directly from the node-labelled Gubbins^47^ phylogeny. For each isolate within a DCC, TBL was defined as the length of its terminal branch, whereas TRL was defined as the cumulative branch length from the corresponding DCC ancestral node to the isolate.

### Core- and accessory-genome phylogenetic reconstruction

To compare the evolutionary structure encoded by the core and accessory genomes, two complementary phylogenetic trees were reconstructed using the diversity-preserving representative isolate set. The core-genome tree was inferred from the concatenated alignment of the 3,001 quality-controlled stable core genes. For the accessory genome, gene presence–absence profiles were extracted from the Panaroo^27^ output. Genes assigned to the stable core genome were removed, and the remaining accessory-gene clusters were represented as a binary matrix, with 1 indicating presence and 0 indicating absence in each isolate. A distance matrix was calculated from these binary profiles and used to reconstruct an accessory-genome phylogeny. The correspondence between the core- and accessory-genome trees was evaluated by comparing the placement and internal organization of the seven DCCs. Phylogeny trees were visualized in *FigTree* (v1.4.4)^48^ or *iTOL*^*49*^.

### Identification of stem-branch mutations and estimation of *r/m*

DCC ancestral nodes were identified from the node-labelled, recombination-aware phylogeny generated by Gubbins^47^. For each DCC, the stem branch was defined as the branch connecting its immediate ancestral node to the most recent common ancestor of the DCC. Branch-specific substitutions and recombinant regions were extracted directly from the Gubbins^47^ ancestral reconstruction and recombination output. Individual SNPs were assigned to genes according to their coordinates in the ATCC 19977 reference genome. SNPs located within Gubbins-predicted recombinant regions were classified as recombination-derived SNPs (*r*), whereas substitutions outside these regions were classified as mutation-derived SNPs (*m*). For each gene and DCC stem branch, the recombination-to-mutation ratio was calculated as:

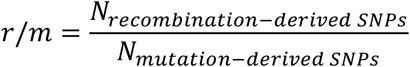

Genes without mutation-derived SNPs were reported as having an undefined r/m value rather than being assigned an arbitrary ratio. For pooled analyses, the numbers of recombination- and mutation-derived SNPs were first summed across DCC1–DCC7, and the overall *r/m* ratio was subsequently calculated from the aggregated counts.

To characterize evolutionary changes after DCC emergence, the same procedure was applied to all internal and terminal branches descending from each DCC ancestral node. Branches or isolates identified as extreme recombination outliers during quality control were excluded before aggregation (Supplementary Fig. 3B, C). Gene-level *r/m* values on the DCC stem branches were then compared with those accumulated across the corresponding post-emergence branches to identify changes in the relative contributions of homologous recombination and de novo mutation.

### Mutational events and selective pressure

Branch-specific mutational events and gene-level selective pressures were obtained using our previously established framework^29^. Briefly, non-recombinant mutations on the stem branch leading to each DCC were classified as pre-expansion events, whereas mutations on descendant branches were classified as post-expansion events. Coding mutations were classified as synonymous or nonsynonymous, and gene-level pN/pS ratios were corrected for synonymous and nonsynonymous mutational opportunities using the observed nucleotide substitution spectrum. Genes showing high-confidence shifts in selective regime between the two evolutionary phases were retained for comparison with changes in r/m in the present study.

### Identification of selective-pressure switches

Because the numbers of mutation events were limited for many individual genes, changes in selective pressure between the pre- and post-expansion phases were quantified using a Bayesian posterior framework. For each gene and evolutionary phase, the probability *θ* that an observed mutation was nonsynonymous was modelled using a binomial likelihood and a uniform Beta(1,1) prior:

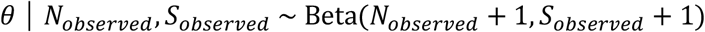

Each posterior draw of *θ* was converted to a mutation-spectrum-adjusted selection ratio, *ω*, according to:

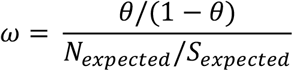

A total of 20,000 posterior draws were generated independently for the pre- and post-expansion phases. For each gene, we calculated the posterior median and 95% credible interval of *ω*, the probabilities that *ω*_*pre*_ > 1 and *ω*_*post*_ > 1, and the probabilities that selective pressure increased or decreased following DCC emergence.

The posterior probability of a purifying-to-positive selection switch was defined as:

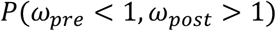

whereas the probability of a positive-to-purifying selection switch was defined as:

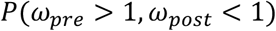

The larger of these two directional probabilities was reported as the posterior switch probability. Genes with a posterior switch probability greater than 0.90 were classified as strong candidates for a shift in selective regime. Gene-level selective-pressure shifts were first evaluated by pooling stem-branch mutation events across DCC1–DCC7 and matching them to post-expansion events obtained from the complete isolate set of each DCC. Analyses were subsequently performed separately for each DCC to determine whether the same genes exhibited convergent shifts in selective pressure across independently expanding lineages.

### Co-expression module identification and functional annotation of selection-shift genes

Co-expression patterns and functional annotations of genes showing shifts in selective pressure were obtained using our previously established framework^29^. Briefly, co-expression modules were defined from normalized expression profiles by pairwise Pearson correlation and hierarchical clustering, and gene functions were assigned by integrating MycoBrowser and eggNOG annotations, structural homology, and manually curated literature.

### Functional enrichment analysis

Genes were functionally annotated using eggNOG-mapper^55^ and mapped to KEGG pathways and modules. Enrichment was assessed separately for genes showing increased or decreased *r/m* after DCC emergence, using all annotated core genes as the background. One-sided Fisher’s exact tests were performed, followed by Benjamini-Hochberg correction. Categories with FDR-adjusted *P* values <0.05 were considered significantly enriched.

## Data Availability

Raw sequencing data are publicly available from the accession numbers listed in Table S1. The custom analysis and visualization scripts supporting this study, together with the phylogenetic trees used in the principal analyses, are available at https://github.com/zhuchendi0520/MAB_core_methods.

