## Supplementary material for "Reduced recombination relative to mutation characterizes dominant circulating clones of *Mycobacterium abscessus*": FigureS1-S6

Chendi Zhu, et al.

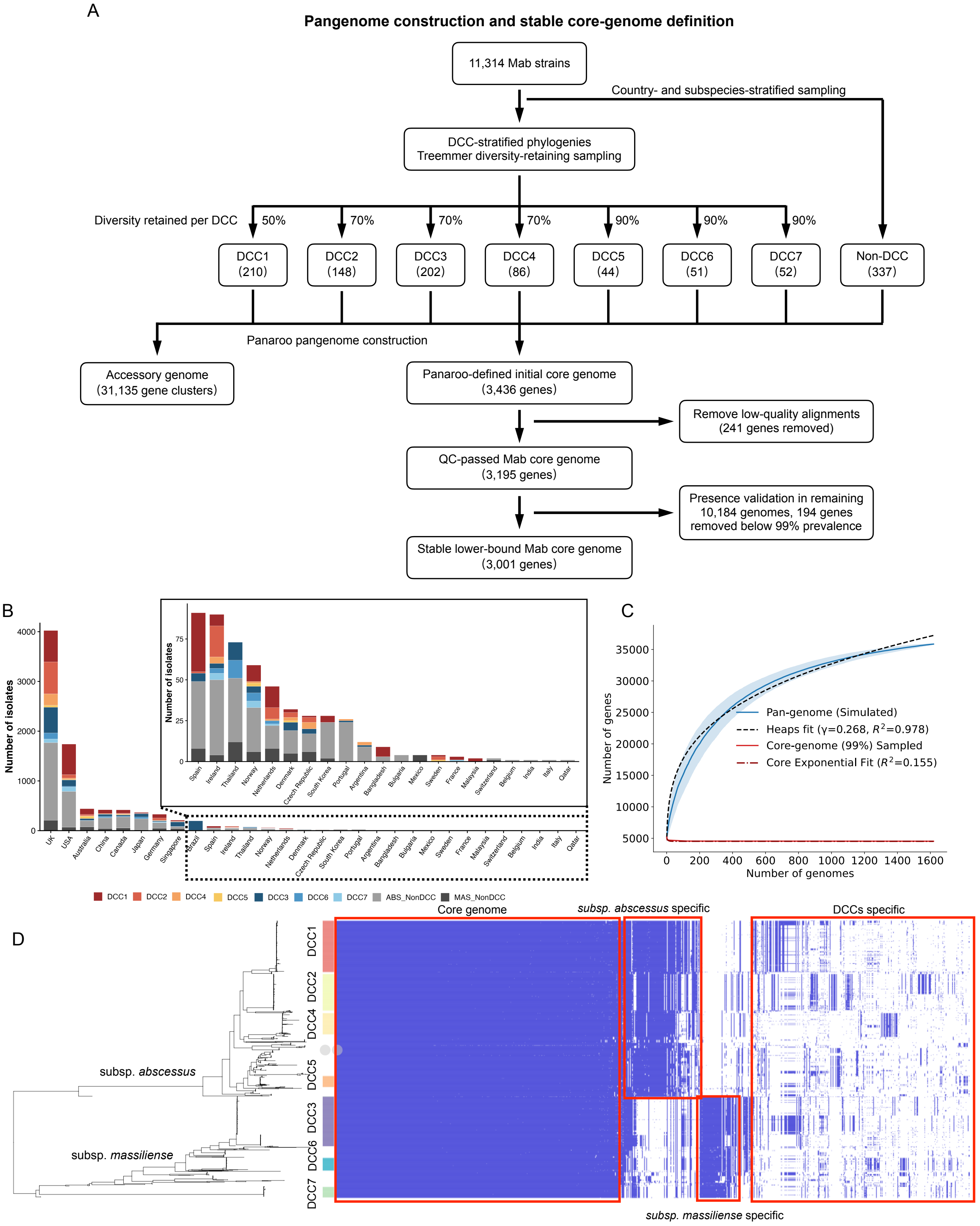


**Supplementary Figure 1. Diversity-preserving sampling and validation of the stable *Mycobacterium abscessus* core genome.** **(A)** Workflow for pangenome construction and core-genome definition. From 11,314 genomes, DCC isolates were subsampled using Treemmer to preserve phylogenetic diversity, while non-DCC isolates were sampled by country and subspecies, yielding 1,130 genomes for Panaroo analysis. Of 3,436 initially identified core genes, 241 with low-quality alignments and 194 present in <99% of the remaining 10,184 genomes were excluded, yielding a stable lower-bound core genome of 3,001 genes. **(B)** Geographic composition of *abscessus* and *massiliense* isolates stratified by DCC status. The inset shows countries with smaller sample sizes. **(C)** Pangenome and core-genome accumulation curves for DCC1. Heaps' law fitting (γ = 0.268, $R^2$ = 0.978) indicates an open pangenome, whereas the core genome approaches a plateau. **(D)** Phylogeny-aligned gene presence–absence matrix. Rows represent isolates ordered by the core-genome phylogeny and columns represent gene clusters, showing the stable core genome and subspecies- and DCC-associated accessory-gene patterns.

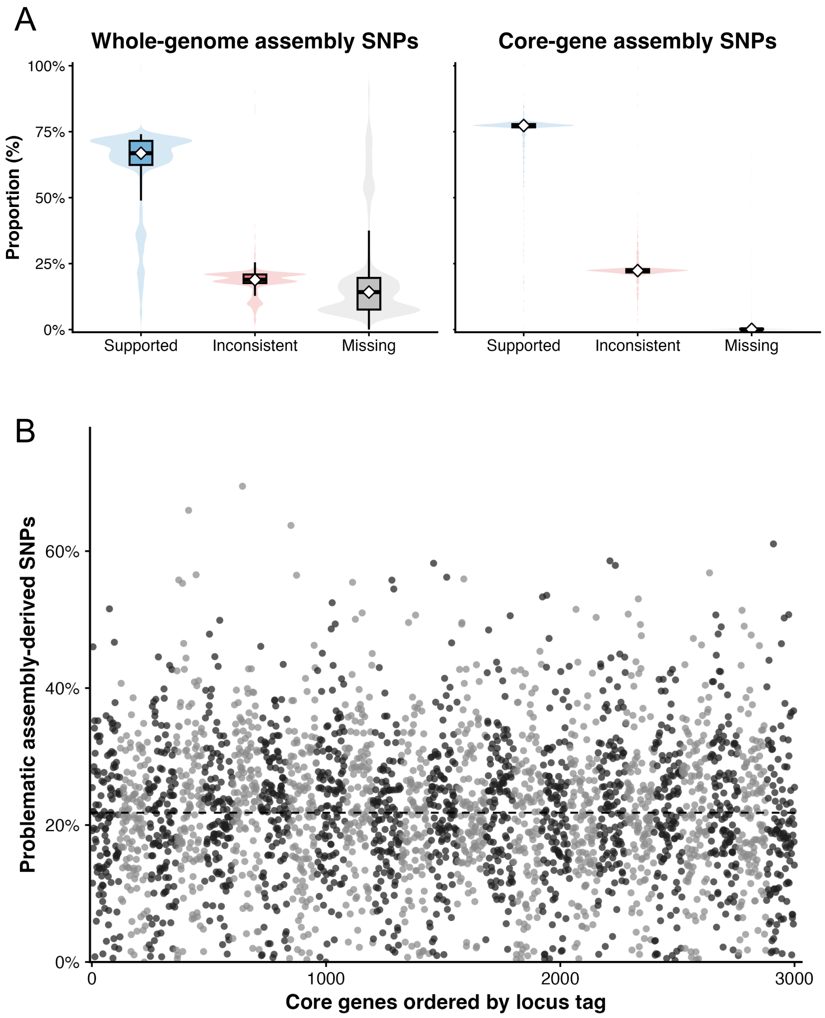


**Supplementary Figure 2. Evaluation of assembly-derived SNP concordance with direct read mapping. (A)** Distribution of supported, inconsistent and missing assembly-derived SNPs across individual isolates, shown for the whole genome and stable core genome. Violin plots show distributions across isolates, with boxplots indicating medians and interquartile ranges. **(B)** Gene-level distribution of inconsistent or missing assembly-derived SNPs across the stable core genome. Core genes are ordered by locus tag, and each point represents the proportion of problematic SNPs within a gene. The dashed line indicates the genome-wide median.

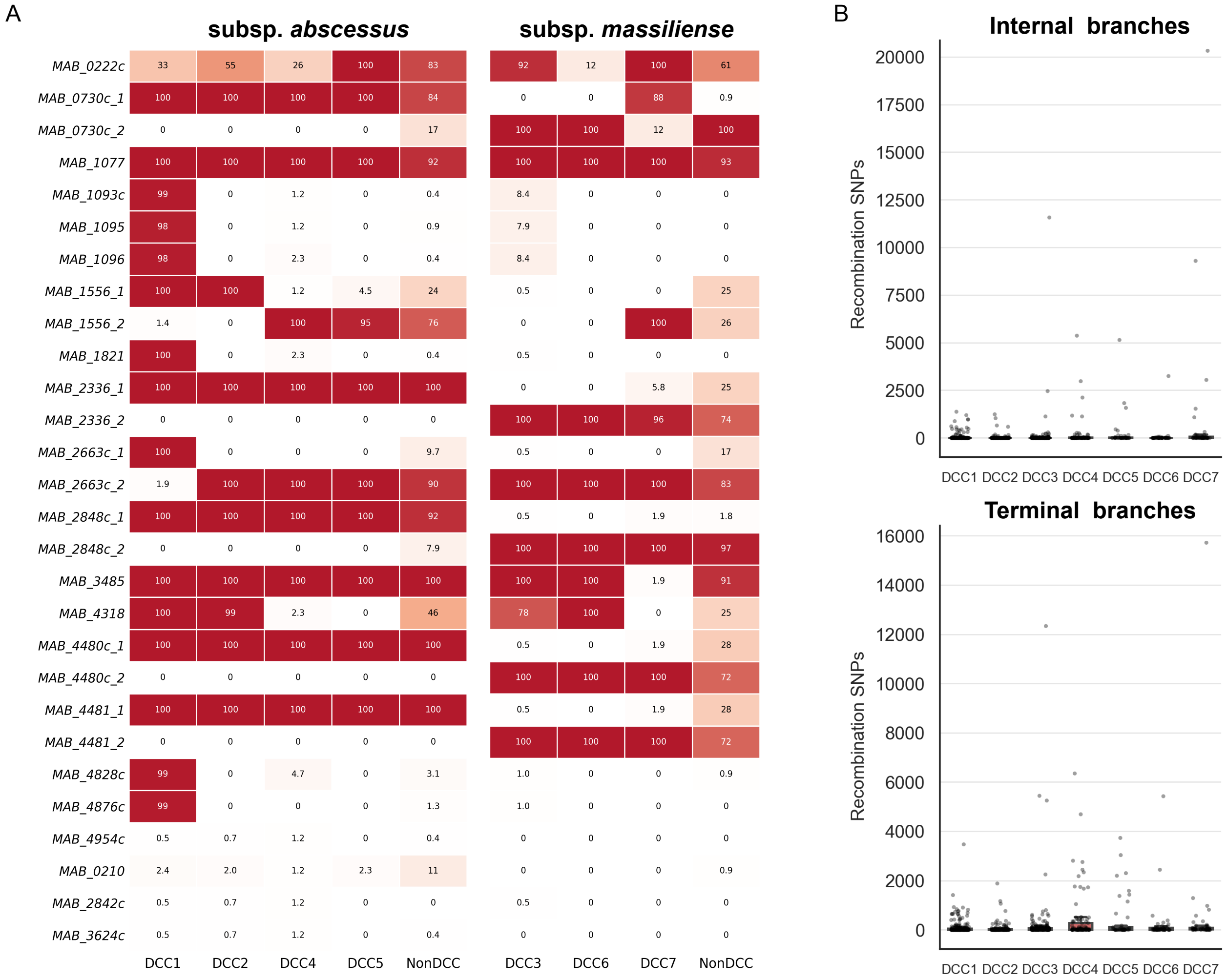


**Supplementary Figure 3. Validation of essential-gene carriage and branch-level recombination patterns across DCCs. (A)** Carriage frequencies of previously reported essential genes not consistently classified within the stable core genome. Frequencies are shown separately for *M. abscessus* subsp. *abscessus* and subsp. *massiliense*, with DCCs compared with their corresponding non-DCC populations. Values indicate the percentage of isolates carrying each Panaroo gene cluster. Multiple clusters mapping to the same ATCC 19977 locus are shown separately. **(B)** Distribution of recombination-associated SNPs assigned to internal and terminal branches within each DCC. Branches with exceptionally high SNP burdens were evaluated as potential outliers before downstream analyses.

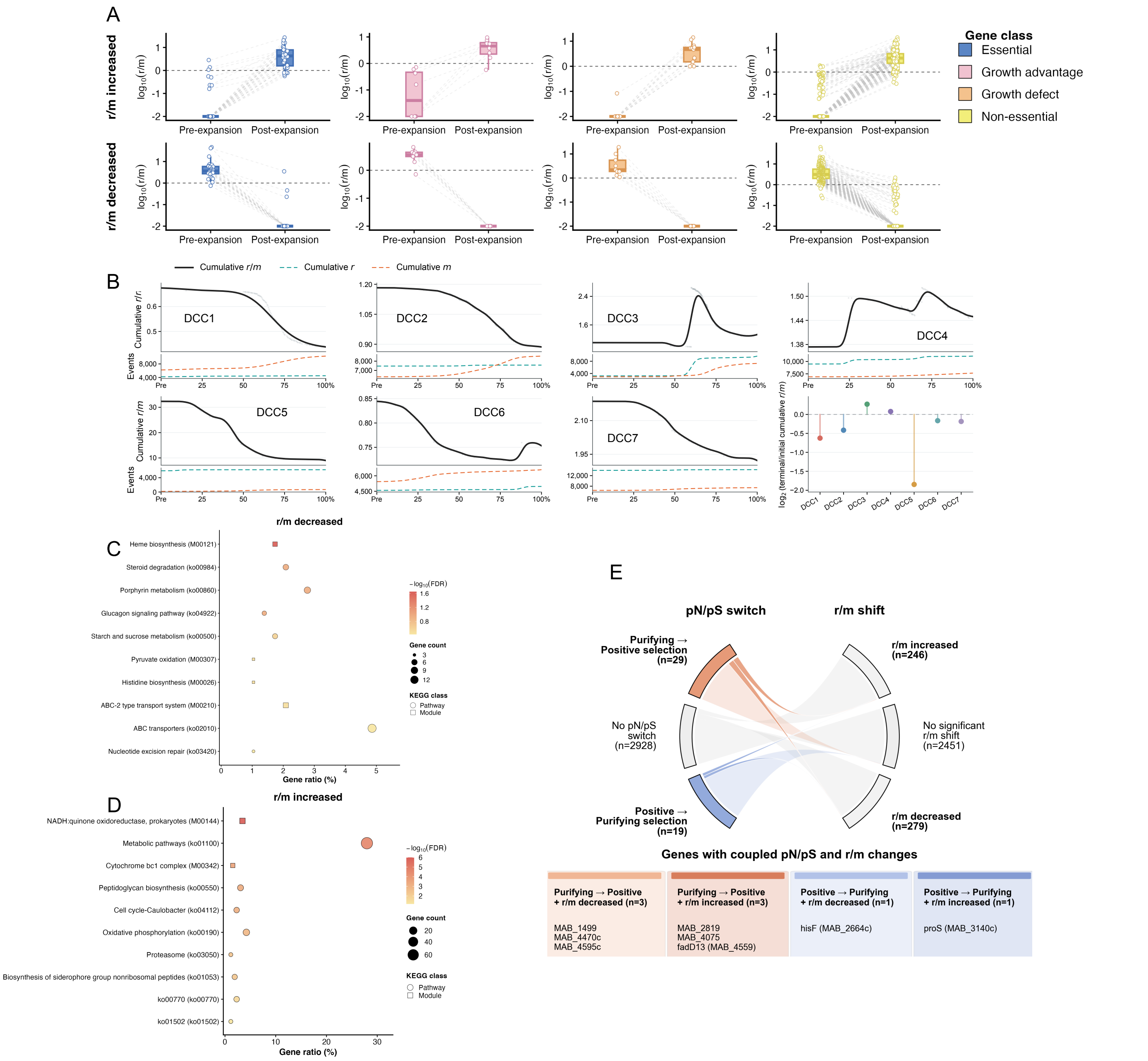


**Supplementary Figure 4. Functional and lineage-specific patterns of evolutionary shifts following DCC expansion. (A)** Paired pre- and post-expansion r/m for genes showing significant shifts, stratified as essential, growth-advantage, growth-defect or non-essential. Zero values were plotted as 0.01 before log10 transformation. Dashed lines indicate r/m = 1. **(B)** Black curves show smoothed cumulative r/m, and teal and orange dashed curves show cumulative recombination-derived and mutation-derived SNP counts, respectively, along normalized tree progression. The lower-right panel shows $\log_2$ changes in terminal versus initial cumulative r/m. **(C-D)** KEGG pathway and module enrichment among genes with significantly decreased **(C)** or increased **(D)** r/m after DCC expansion. Point position indicates gene ratio, point size the number of annotated genes, color −log10(FDR), and shape KEGG pathways or modules. **(E)** Relationship between pN/pS and r/m shifts. Flows connect pN/pS switch categories with increased, decreased or non-significant r/m shifts. The accompanying table shows genes with shifts in both metrics.

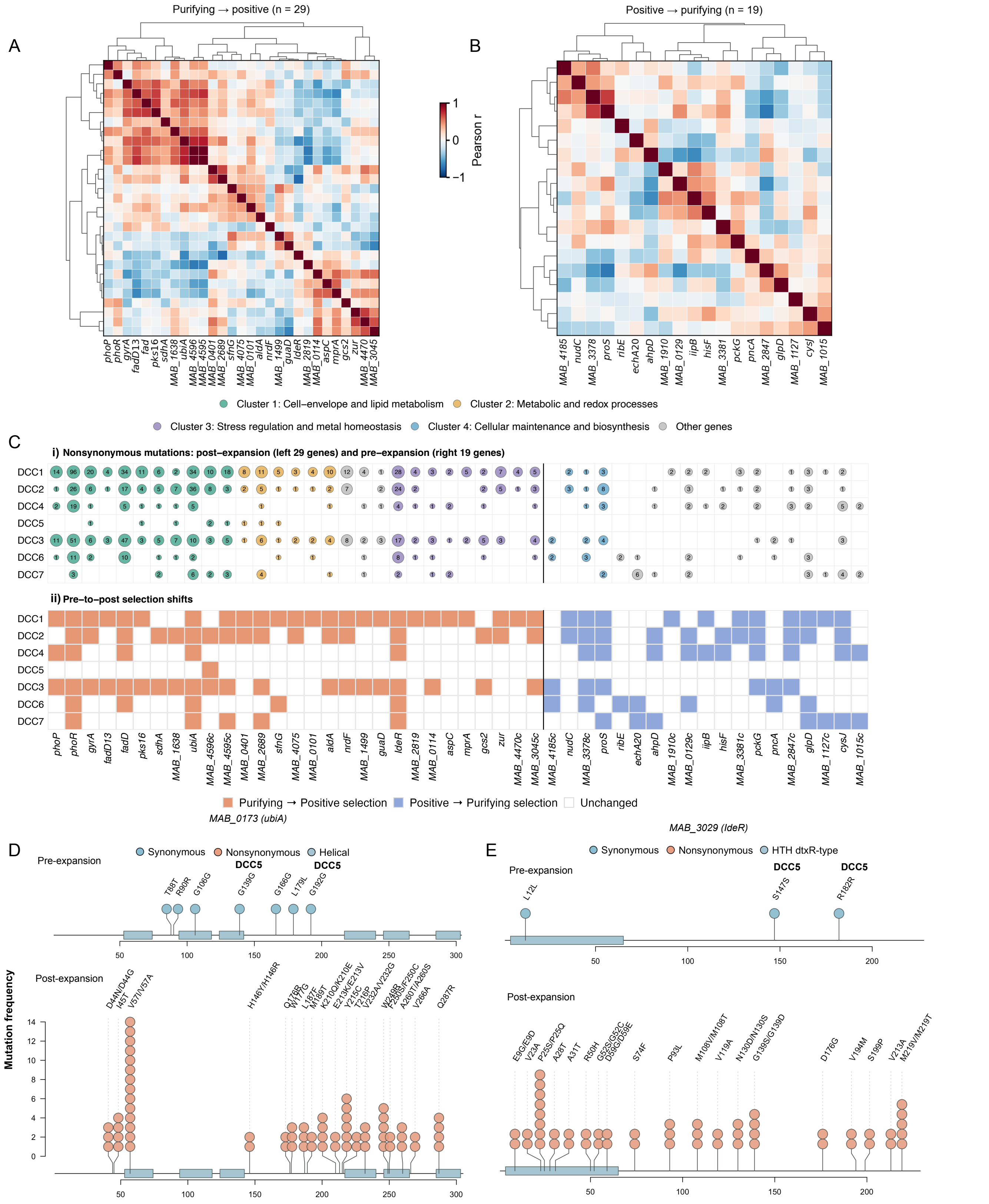


**Supplementary Figure 5. Stem-branch mutation patterns across DCCs. (A–B)** Co-expression structure of genes showing high-confidence shifts in selective pressure. Pairwise Pearson correlations were calculated from batch-corrected expression profiles and hierarchically clustered separately for genes shifting from purifying to positive selection **(A)** and from positive to purifying selection **(B)**. Major co-expression modules and representative regulatory and functional genes are indicated.**(C)** Distribution of high-confidence selective-shift genes across individual DCCs and their associated functional modules. **i)** Although the specific genes affected differed among DCCs, recurrent changes converged on shared biological processes, including cell-envelope remodeling, lipid metabolism, stress responses, metal homeostasis, and cellular maintenance. **ii)** DCC-specific occurrence of high-confidence pN/pS switch genes identified in the pooled analysis. Orange indicates purifying-to-positive shifts, blue indicates positive-to-purifying shifts and white indicates no switch across pN/pS = 1. **(D–E)** Protein-level distribution of mutation-derived SNPs in representative genes with recurrent purifying-to-positive selection shifts: *ubiA* (*MAB_0173*; **D)** and *IdeR* (*MAB_3029*; **E)**. Upper and lower tracks show mutations on DCC stem branches and descendant branches after expansion, respectively. Circle height indicates mutation frequency; blocks denote protein domains.

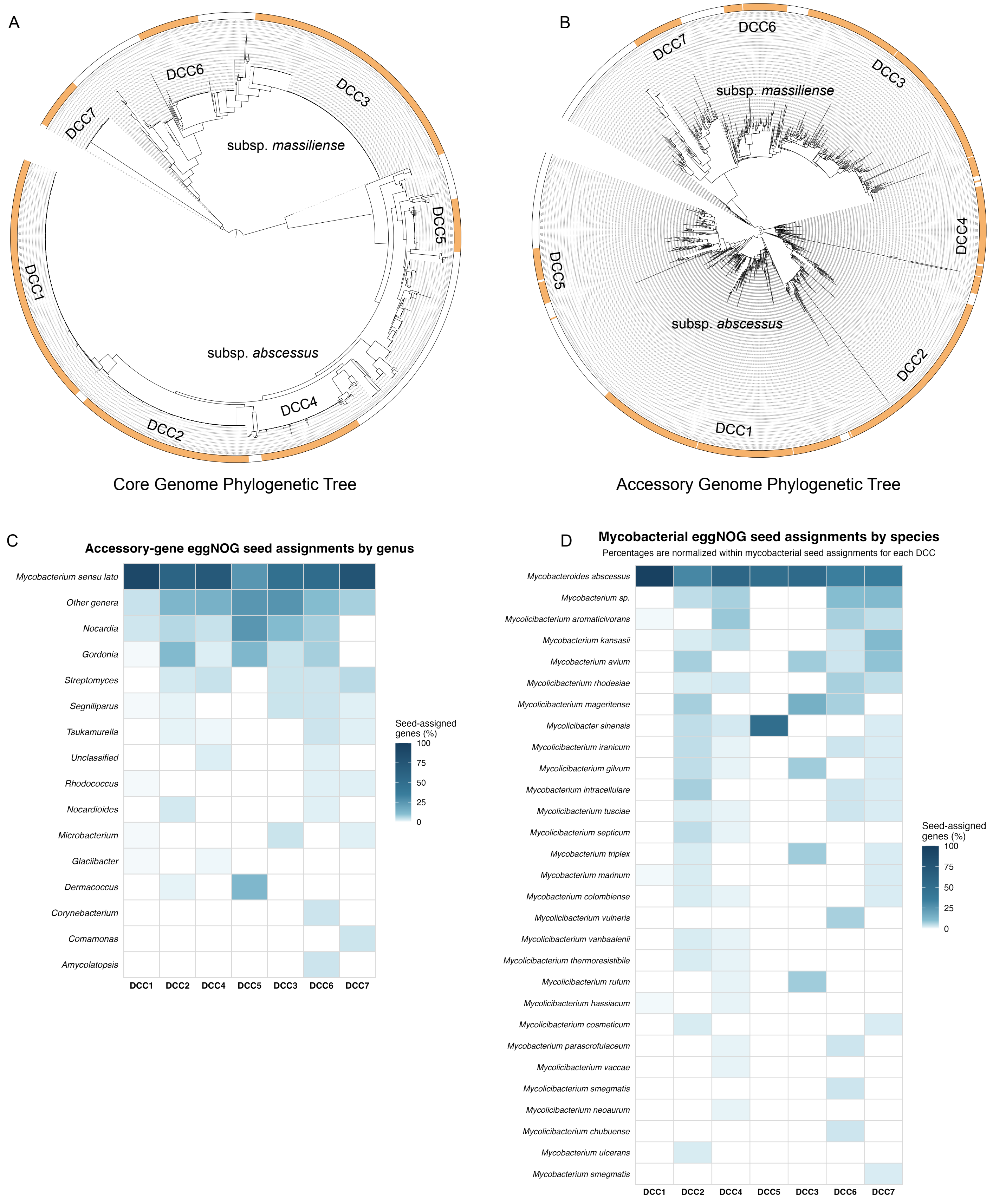


**Supplementary Figure 6. Population structure and taxonomic affinities of DCC-associated accessory genes. (A-B)** Circular phylogenies reconstructed from the stable core-genome alignment **(A)** and accessory-gene presence–absence matrix **(B).** Orange outer segments denote DCC isolates. Both reconstructions recover the major subspecies division and broadly concordant DCC structures. **(C)** Genus-level taxonomic affinities of eggNOG seed sequences assigned to DCC-associated accessory genes. Members of Mycobacterium sensu lato were grouped into a single category. Values are normalized within each DCC and shown as percentages of genes with taxonomically assigned seed sequences. **(D)** Species-level composition of mycobacterial eggNOG seed assignments. Values indicate the percentage of mycobacterial seed-assigned accessory genes attributed to each species within each DCC.
